# Optimizing Protocols for MicroRNA Profiling of Infant and Toddler Stool

**DOI:** 10.1101/2025.04.01.646630

**Authors:** David A. Armstrong, Shannon M. Soucy, Meghan E. Muse, Fred W. Kolling, Heidi W. Trask, Alexandra L. Howell, Hannah E Laue, Anne G. Hoen, Jiang Gui, Brock C. Christensen, Juliette C. Madan, Margaret R. Karagas, Caitlin G. Howe

## Abstract

**Background:** MicroRNAs (miRNAs) are increasingly being investigated as potential biomarkers for child development and disease. Although a growing number of studies are utilizing infant and toddler stool for transcriptomic analyses, no studies have compared protocols for preserving and extracting miRNAs from this specimen type, despite unique challenges, including abundant levels of RNAses and microbial RNA.

**Methods:** To address this, we first compared three commercially available kits and four preservation methods for their ability to yield high quality RNA from infant and toddler stool (Phase 1). RNA quality was determined by fragment analyzer.

**Results:** Of the three RNA extraction kits compared, Zymo BIOMICs yielded the highest overall RNA Quality Number (RQN) (median (range) RQN 9.4 (5.7-10.0)). Of the four preservation methods tested, stool collected in RNAlater and Zymo DNA/RNA Shield Fecal Collection Tubes yielded the highest two RQNs (median (range) RQN 9.8 (5.7-10.0) and 9.4 (5.4-10.0), respectively), which did not differ significantly from each other (*p* = 0.47). Second, using miRNA-seq we directly compared miRNA profiles for RNA extracted using the Zymo BIOMICs kit from paired aliquots of the same stool sample from four infants collected into RNAlater and Zymo DNA/RNA Shield Fecal Collection Tubes (Phase 2). Given that microbial sequences greatly outnumber human miRNAs in stool, reads were first classified as human versus microbial prior to aligning human-classified reads to miRBase v22.1. The percentage of reads classified as human and the percentage of human reads aligning to miRBase did not differ for samples collected in RNAlater versus Zymo Shield (*p* = 0.12 and *p* = 0.86, respectively). Furthermore, after multiple testing correction, normalized miRNA counts did not differ significantly between the two preservatives for any of the 42 human miRNAs detected across the eight samples.

**Conclusions:** Collecting infant and toddler stool in either RNAlater or Zymo DNA/RNA Shield Fecal Collection Tubes, when paired with RNA extraction using the Zymo BIOMICs extraction kit, yielded high-quality RNA with similar human miRNA profiles. Moreover, of the 42 miRNAs that were detected, several (i.e., miR-194a-3p, miR-200c-3p, miR-26a-5p) are thought to contribute to overall gut homeostasis. These findings may inform protocols for future studies that aim to profile miRNAs in infant and toddler stool to evaluate their potential utility as biomarkers for children’s health.

## Background

MicroRNAs (miRNAs) are a class of small non-coding RNAs, 19 to 24 nucleotides in length, that regulate a range of developmental and physiological processes by modulating gene expression through post-transcriptional RNA interference and other gene silencing pathways [1, 2]. MiRNAs have been shown to exert their influence by regulating dosage-sensitive genes for which small fluctuations in protein expression may have significant functional consequences [3]. Moreover, a single miRNA can target multiple genes and conversely, a single gene can be regulated by several different miRNAs [4]. As of the latest version of miRBase (version 22) 2,637 annotated human miRNA loci have been identified [5]. Collectively, these miRNAs have a profound impact on gene expression and are estimated to regulate up to 60% of protein-coding genes.

Increasingly, miRNAs have gained attention for their potential as diagnostic tools, and dysregulation of miRNAs has been implicated in various diseases and pediatric outcomes. For example, many studies have profiled miRNAs in diverse specimens with the goal of identifying potential biomarkers for pediatric cancers, such as neuroblastoma and hematological cancers [6], as well as infectious diseases [7] and immune disorders [8, 9]. MiRNAs also play important roles in maintaining a healthy pregnancy, and their levels in the placenta, maternal circulation, and cord blood have been associated with a number of pregnancy and birth outcomes, including gestational diabetes, preeclampsia, low birth weight, and gestational duration, further illustrating their potential to serve as early biomarkers of child health [10–21].

Despite its utility for studying important aspects of children’s health, including the developing microbiome, stool from infants and toddlers has not been used widely for studying miRNAs. Notably, more than 50 published studies have utilized adult stool for RNA extraction, but only nine published studies have focused on RNA from stool of young children [22–30]. Currently, several challenges exist for accurately profiling miRNAs in stool, such as the presence of components that inhibit downstream methodologies (e.g., bilirubin), which can make it difficult to isolate high-quality RNA from stool or to conduct downstream sequencing [31]. Another major challenge is the high prevalence of both host and microbiome-derived RNases resulting in RNA degradation [32, 33]. For transcriptomic applications, it has therefore been recommended that fixatives blocking *de novo* transcription and RNA decay be used for stool collection to preserve the RNA composition of the sample [34]. Differences in specimen handling can further compound this issue. This is a particularly important consideration when focusing on stool from infants and young toddlers, given that most children of this age are still using diapers, which may lead to substantial variability in the time between sample void and collection. RNA extraction efficiency from stool can also differ by kit, influencing RNA quality and downstream transcriptomic analyses.

Although a prior study by Reck, *et al.* [35] compared commercially available RNA preservation solutions and extraction kits for stool, it focused exclusively on samples from adults with the goal of informing best practices for metatranscriptomics. As of yet, there has been no comparison of preservatives or extraction kits for isolating RNA from stool of infants and young toddlers for the detection of human transcripts. While several recent studies have aimed to profile human miRNAs in stool from this early life stage, methods for stool collection and RNA extraction varied. Additionally, most studies did not report using any RNA stabilizing solution at the time of stool collection [22–30]. Thus, to inform best practices for future studies, the primary objectives of this study were to (1) compare RNA quality of stool collected from infants and young toddlers (<2 years of age) in different preservative conditions, focusing on protocols used in prior published studies of either child or adult stool [24, 36–41] and, (2) compare the quality of RNA extracted from stool collected from infants of young toddlers using different commercially available and widely used stool RNA extraction kits. For protocols yielding high-quality RNA, we additionally aimed to (1) compare miRNA profiles across protocols and (2) identify the most abundant human miRNAs present in stool from infants and young toddlers to serve as a resource for future studies on this topic.

## Methods

### Sample collection

All study protocols were approved by the Dartmouth College Committee for the Protection of Human Subjects, which is the Institutional Review Board at Dartmouth College. Parents of participating children provided written informed consent, and stool samples and their associated data were de-identified.

In the first phase of the study, a total of six parent-child pairs were enrolled (**Figure 1**).

**Figure 1.**
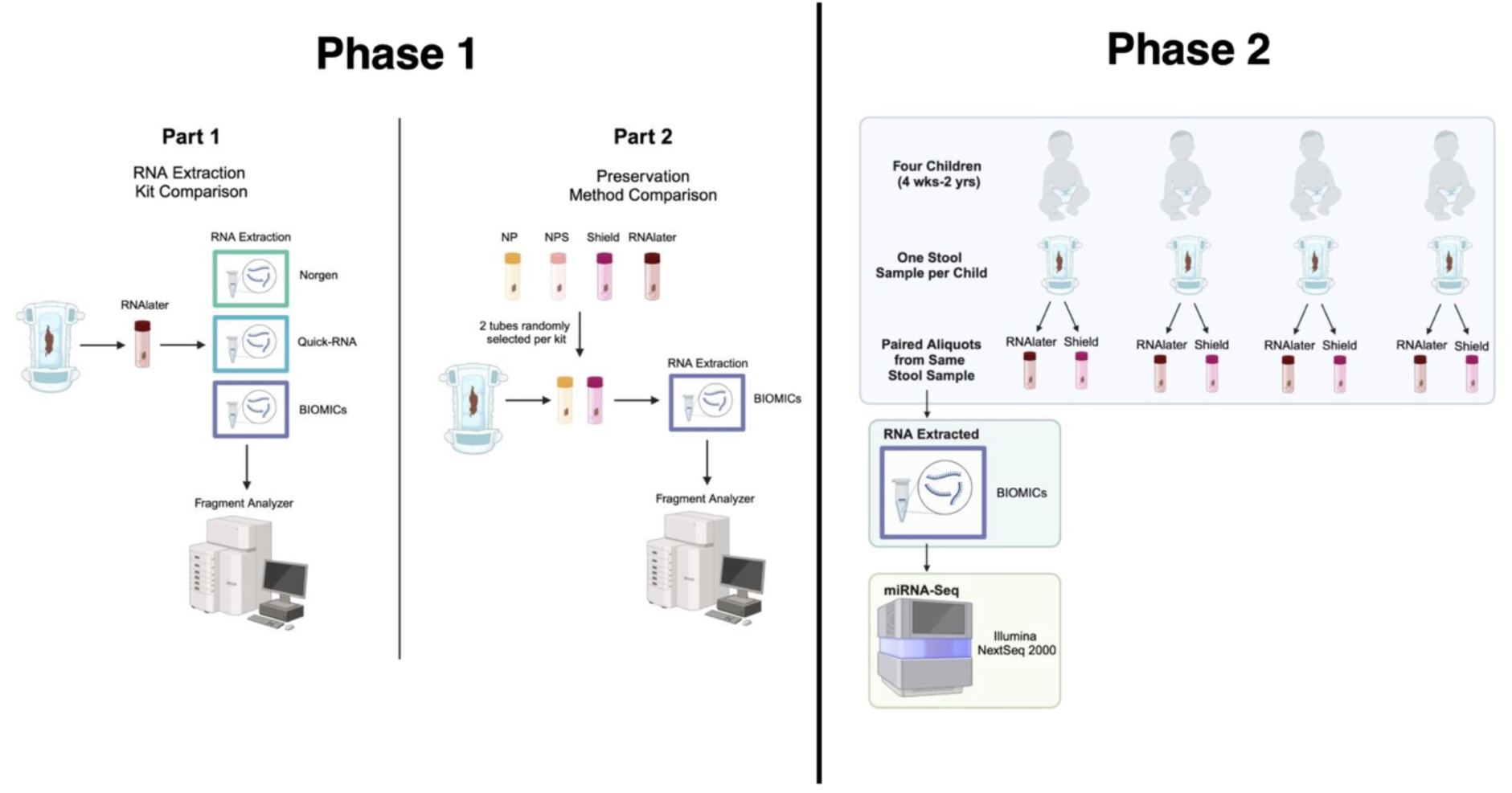
Study Design. <u>Phase 1</u> of the study consisted of two steps: (1) comparing RNA quality when using three different RNA extraction kits for stool collected in RNAlater and (2) comparing RNA quality for RNA extracted from stool collected in four different preservation conditions, including no preservative (NP), with RNA extracted using the BIOMICs RNA extraction kit. For each step, RNA quality number was determined using a fragment analyzer. <u>Phase 2</u> of the study entailed collecting paired aliquots from the same stool sample into RNAlater versus Shield for four infants. RNA was extracted using the BIOMICs RNA extraction kit. MiRNAs were then quantified for each sample using miRNA-seq. (Figure created in Biorender).

In the second phase of the study, a total of four parent-child pairs were enrolled (**Figure 1**). Inclusion criteria were the same for each phase of the study: children needed to be between 4 weeks and 2 years of age and using diapers. For both phases of the study, participants were recruited by word-of-mouth and flyer advertisements at Dartmouth College and local daycares. In the first phase of the study, the parent was told that they could participate up to six times and receive a $25 gift card each time they participated. Upon enrollment, participants received up to six collection kits (depending on the number of times that they decided to participate), containing two of any of the following collection tube types: Stool Nucleic Acid Collection and Preservation Tubes (Norgen, Thorold, ON, Cat 45660) (Norgen), DNA/RNA Shield Fecal Collection Tubes (Zymo, Irvine, CA Cat R1011) (Shield), a Sarstdedt feces tube (Sarsdedt, Newton, NC, Cat 80.734.001) containing 9 mL of RNAlater (ThermoFisher Scientific, Waltham, MA Cat AM7020), or an empty Sarstdedt feces tube (no preservative). Tube types were randomized to each kit prior to distribution. For the second phase of the study, to ensure representation of different children, parents were told that they could participate once and received a $25 gift card for participating. During this second phase, each participant received one stool collection kit, which included the two tube types yielding the two highest median RQN values during the first phase of the study.

Each stool collection kit included a specimen form and detailed written instructions for collecting the samples. As soon as the parent noticed that their child had a bowel movement, they were instructed to transfer a nickel-diameter (“marble-sized”) mass of stool from the diaper into each of the provided tubes. These tubes were cylindrical screw-cap fecal collection containers with an attached spoon to assist with collection. The parent was then instructed to immediately cap the tube, place it in a provided biohazard bag, and store the specimen(s) in their home freezer until collection by the study team. If the tube contained a preservative, the participant was instructed to first invert the tube multiple times to ensure that the sample was completely immersed in the preservation fluid. During the first phase of the study, parents were told that for each tube they could collect stool from the same diaper or from different diapers. During the second phase of the study, parents were instructed to collect two stool aliquots from the same diaper into each of the two provided tubes. In addition to collecting stool from their child, parents were asked to complete the following information on the specimen form: the date and time of stool collection, the child’s age (in months), the child’s biological sex assigned at birth, whether the child was currently breastfed, and the brand and type of diaper used for stool collection. Although parents were not advised to use a specific type of diaper to facilitate collection, all participating parents reported using disposable diapers. Participants were instructed to coordinate transfer of the samples to the laboratory within 48 hours of collection. Samples were transported by study staff on cold packs to the COBRE Center for Molecular Epidemiology Laboratory at Dartmouth College. Upon arrival to the laboratory, samples were immediately transferred to a -80°C freezer where they were stored until processing. The median (range) duration that the samples were stored in the participant’s home freezer prior to being transferred to the laboratory’s -80°C freezer was 46 (1-142) hours for Phase 1 of the study, and 25 (19-86) hours for Phase 2 of the study. The median (range) duration that samples were on cold packs during transport to the laboratory was 52 (23-99) minutes for Phase 1 of the study, and 23 (10-80) minutes for Phase 2 of the study. *RNA Extraction Kits, Extraction Protocols and RNA Quantitation*.

Three different stool RNA extraction kits were tested for their ability to isolate high quality RNA from infant/toddler stool: Stool Total RNA Purification Kit (Norgen Biotek 49500), (2) Quick-RNA Fecal/Soil Microbe Microprep Kit (Zymo R2040) (Quick-RNA) and (3) Zymo BIOMICs RNA Miniprep (Zymo R2001) (BIOMICs). These kits were selected based on the following criteria: [42] recent publications had reported using the kit for RNA isolation from either infant/toddler or adult stool [23, 41], (2) there was current market availability, as some kits referenced in prior literature were no longer available when the study was initiated [43], (3) the protocol involved a straightforward column-based format, and (4) the protocol did not include a phenol/chloroform extraction step. The latter two selection criteria were based on technical and safety considerations. Given prior work from Reck, *et al.* (2015), which identified RNAlater as the optimal stool RNA preservation solution for adult stool (based on RNA Integrity Number), infant/stool samples collected in RNAlater were used for this initial kit comparison. Approximately 100-200 milligrams of stool were used for each extraction based on manufacturer recommendations. For the Norgen and Quick-RNA kits, RNA was extracted following the manufacturer’s written instructions without any modifications. However, the BIOMICs kit protocol was modified in accordance with the manufacturer’s technical support guidelines. Specifically, Step 4 of the <u>Sample Preparation</u> section of the manufacturer’s protocol was modified by increasing the ratio of RNA Lysis Buffer to supernatant from 1:1 to 3.6:1 (i.e., 1250 µl Lysis Buffer: 350 µl of sample supernatant). Additionally, within the <u>Total RNA Purification</u> section of the protocol (Step 1), the volume of 100% ethanol was adjusted to 1600 µl to account for the increased Lysis Buffer input. These modifications were implemented to reduce sample precipitate and improve RNA yields. All other steps were followed according to the manufacturer’s instructions. RNA was quantified by Qubit (ThermoFisher, Waltham, MA) and integrity (RQN) was measured using a fragment analyzer (Agilent Technologies, Santa Clara, CA) by the Genomics and Molecular Biology Shared Resource at the Dartmouth Cancer Center (Lebanon, NH).

### miRNA Sequencing

MiRNA-sequencing (miRNA-seq) was conducted by the Genomics and Molecular Biology Shared Resource at the Dartmouth Cancer Center (RRID: SCR-021293) for the four sets of paired samples collected in phase 2 of the study (*n*=8 samples total). After RNA quantification by qubit and integrity measurement using a Fragment Analyzer instrument (Agilent), 100 ng of total RNA was used as input into the Qiaseq smRNA library preparation kit (Qiagen) and processed following the manufacturer’s instructions, using 16 cycles for final library PCR. Samples were uniquely indexed and pooled for sequencing on a NextSeq 2000 instrument (Illumina), targeting 10 million reads per sample.

### Data Pre-Processing

Data pre-processing was conducted by the Dartmouth Center for Quantitative Biology Genomic Data Sciences Core. Raw reads were processed with UMItools v1.1.5 [44] to remove umi barcodes. Reads were then classified using centrifuger v.1.0.4-r153 with a custom database built from refseq bacterial, archaeal, and viral genomes plus the representative human genome (GRCh38.p14) to distinguish human reads from microbial reads. Reads were further classified as human *versus* microbial using kneaddata v.0.12.0, with reads mapping to GRCh38 identified as human. Reads that were assigned as human by both centrifuger and kneaddata were retained as the final confident set of human reads for downstream analyses.

All miRNA sequences were downloaded from release 22.1 of miRbase. The file was filtered for unique human miRNAs and any uracil bases were converted to thymine bases. The accompanying gff file was used to ascertain the genome coordinates for each sequence. To account for any transcriptional wobble and increasing mapping rates, an additional five base pairs of sequence were added to each end of the miRNA sequence in the reference file. The confident set of human reads identified using centrifuger and kneaddata were mapped against this padded reference using bowtie2 v2.4.4 with custom settings that increased the probability of seed matching to short miRNA reference sequences, reduced the size of the seed, reduced the size of the window for extending the seed, and enabled mismatches within the seed (-D 20 -R 3 -N 1 -L 12 -i S,1,0.50). The mapped reads were filtered with samtools v1.11 such that matches were required to be between 16 and 28 base pairs in length, with no gaps, fewer than two mismatches, and a MAPQ score greater than one. Reads were then deduplicated with UMItools v.1.1.5 before using Samtools v1.11 to quantify reads mapped to each miRNA. Mapped reads were visually inspected with Integrative Genomics Viewer (IGV) [45] v.2.8.7.

### Statistical Analysis

Statistical analyses were conducted in R (v.4.3.1). The median and interquartile range values for RQN were calculated for each kit and preservation method. Kruskal-Wallis tests were used to assess where RQNs differed by kit and preservation method. If test results were statistically significant (*p* < 0.05), pairwise Wilcoxon tests were then conducted to identify which pairs of kits or which pairs of preservation methods differed.

MiRNA counts were normalized using EdgeR (v.3.42.4) with the Trimmed Mean of M-values (TMM) method to remove technical variation related to variability in library sizes. The pheatmap (v.1.0.12) package was used to explore how samples clustered [46]. MiRNAs were included in the heatmap if at least one sample had a minimum of 10 raw read counts. Paired t-tests were used to evaluate whether the percent of human reads or the percent of human reads aligning to miRbase differed between the paired stool aliquots collected in the two preservative types yielding the two highest RQNs, and to evaluate whether the normalized counts differed significantly between these preservatives for any of the detectable miRNAs. High confidence predicted target genes of the top 10 most abundant miRNAs were identified using mirDIP (v.5.2) and gene set enrichment analyses for gene ontology (GO) biological processes (2023) and PANTHER pathways (2016) of identified target genes were conducted using enrichR (v.3.2) [47].

## Results

### Participant Demographics

For Phase 1 of the study, a total of 60 stool samples were collected from eight children four to 19 months of age. In Phase 2 of the study, a total of eight stool samples were collected from four children four to 21 months of age. Demographics of the children who participated in each phase are shown in **Table 1**.

**Table 1.** Participant Characteristics.

|  | Phase 1 (*n*=8 infant donors) | Phase 2 (*n*=4 infant donors) |
| --- | --- | --- |
|  | Median (Range) or N (%) | Median (Range) or N (%) |
| Age (Months) | 6.8 (4.0-19.0) | 11.3 (4.0-21.0) |
| Female Sex | 2 (25.0) | 3 (75.0) |
| Breastfed | 4 (50.0) | 3 (75.0) |

### Phase 1: Comparison of Extraction Kits and Preservation Methods

The median RQN for each stool RNA extraction kit is shown in **Figure 2A**.

**Figure 2.**
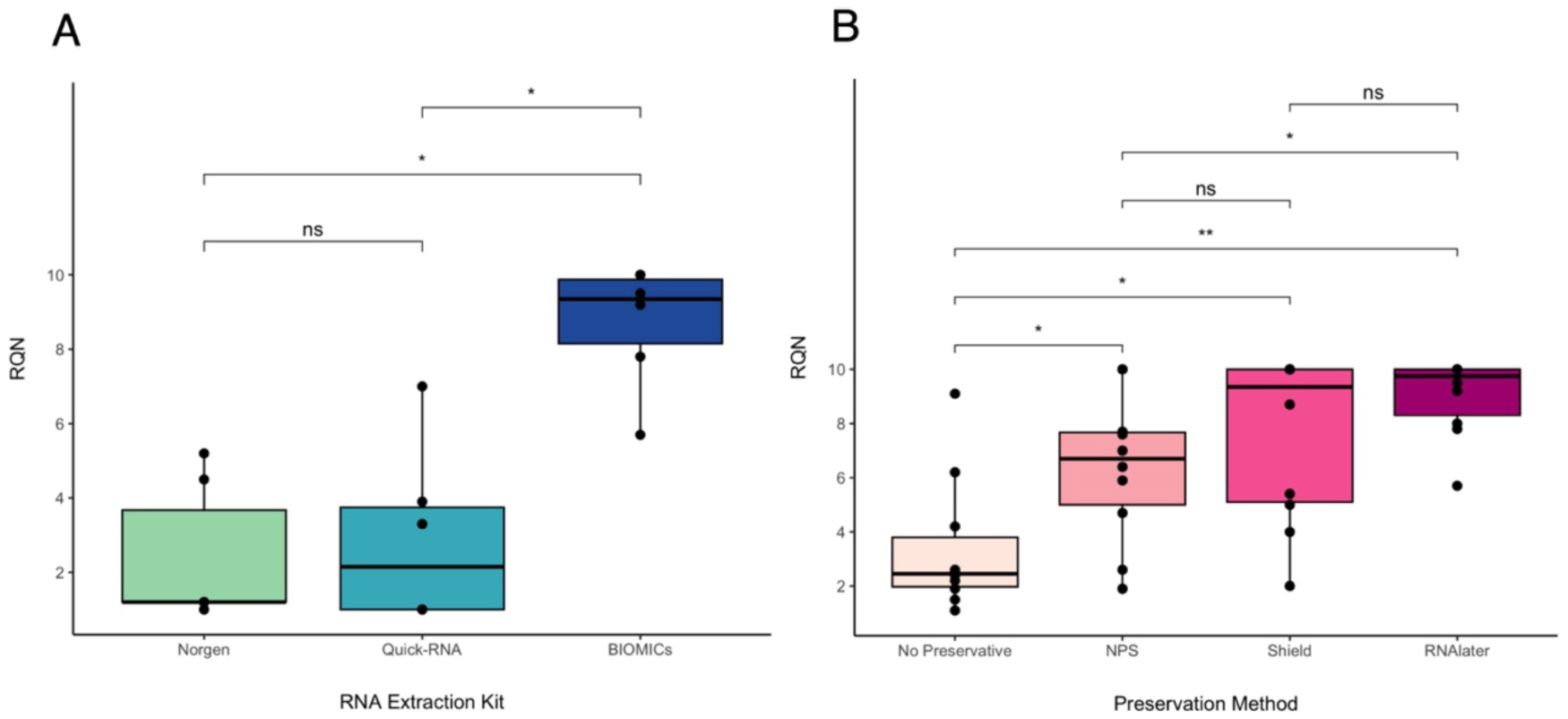
RNA Quality Number by RNA Isolation Kit (A) and Preservative Type (B). RNA was extracted from stool collected in RNAlater using three commercially available kits **(A)**. RNA quality was determined based on RQN using a fragment analyzer. The highest median RQN (9.4) was observed for the BIOMICs extraction kit, followed by the Quick-RNA kit (2.2), followed by the Norgen kit (1.2) (*n*=6 samples per extraction kit). Next, RNA was extracted from samples stored in three different preservative solutions or in the absence of preservative using the BIOMICs kit (*n*=10 samples per collection protocol) (**B**). RNA from stool samples collected in RNAlater had the highest median RQN (9.8), followed by Shield (9.4), followed by NPS (6.7), followed by no preservative (2.5). *P<0.05, **P<0.01. Brackets are not shown if the difference was not statistically significant (P≥0.05)

**Figure 3.**
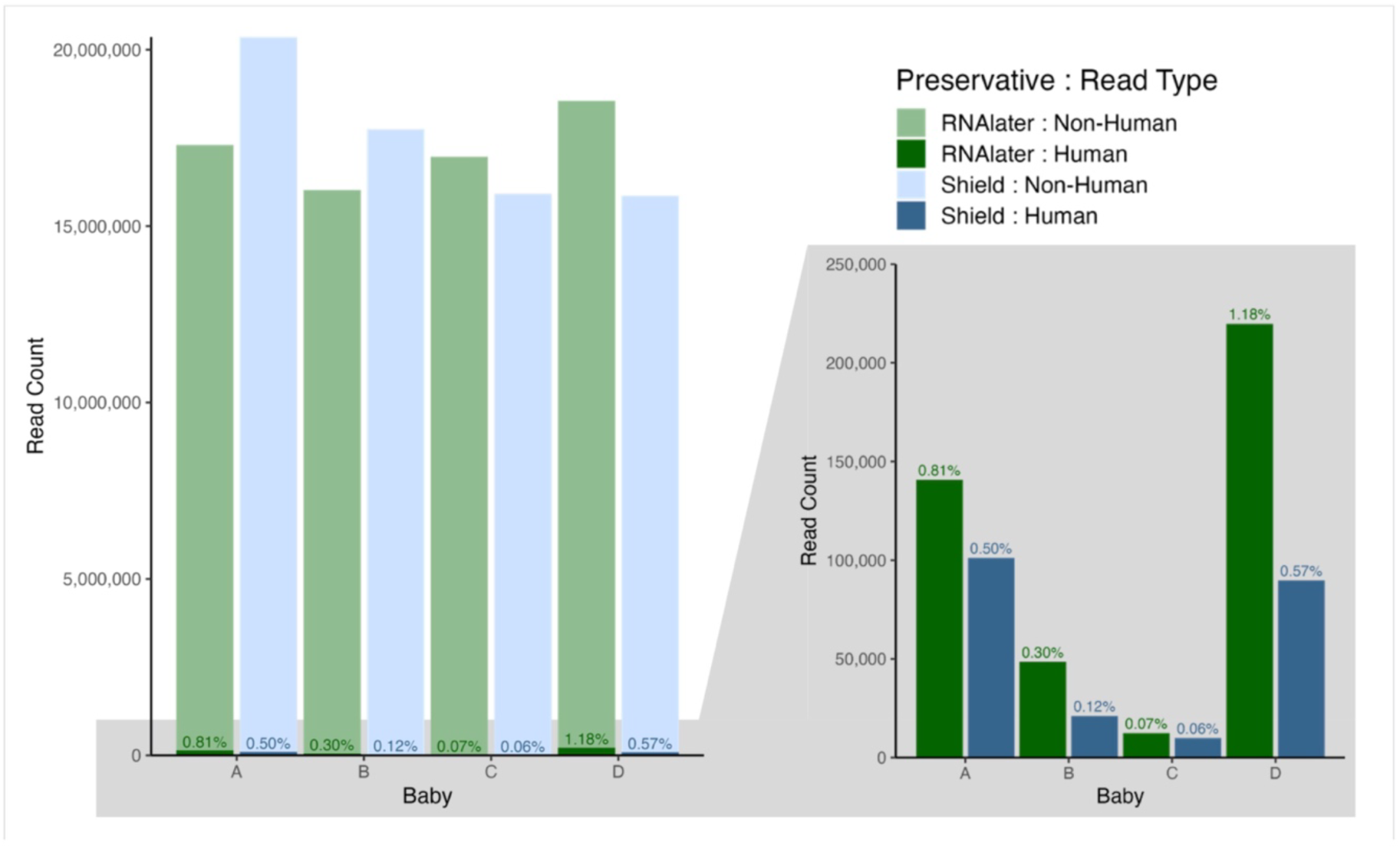
Total Read Counts and Percent Classified as Human. Barplot showing total read counts and the percentage of reads classified as human for paired aliquots of infant stool transferred from four infants’ (A, B, C, D) diapers into RNAlater and Shield preservatives. Percentages reflect the percentage of human reads for the sample.

The BIOMICs kit yielded a higher RQN (median: 9.4, range: 5.7-10) than the Quick-RNA Kit (median: 2.2, range: 1.0-7.0) (*p* = 0.01) or the Norgen kit (median: 1.2, range: 1.0-5.2) (*p* = 0.01). Fragment analysis electropherograms are shown for each stool sample that had the RQN closest to the median RQN for each kit (**Additional File 1**). The median RQN for each preservative type are shown in **Figure 2B**. RNAlater yielded the highest median RQN (9.8, range: 5.7-10.0), which differed significantly from NPS (median: 6.7, range: 1.9-10.0) (*p* = 0.03) and No Preservative (median: 2.5, range: 1.1-9.1) (*p* < 0.01) but not Shield (median: 9.4, range: 2.0-10.0) (*p* = 0.47). Electropherograms are shown in **Additional File 2** for samples with RQNs falling closest to the median value for each preservative condition to illustrate what might reasonably be expected. For all samples, electropherograms included peaks consistent with prokaryotic ribosomal 23, 16 and 5 S subunits with evidence of various degrees of RNA degradation as shown by small RNA peaks/fragments present between 140-1435 nucleotides in length.

### Phase 2: Comparison of miRNA Profiles for Protocols Yielding High Quality RNA

For the eight samples that underwent miRNA-seq, library sizes ranged from 15.9 million to 20.4 million reads. Human reads represented less than 1% of total reads for all but one sample **(Figure3**).

Although samples collected in RNAlater had a higher percentage of human reads (median: 0.56% range: 0.07-1.18%) compared with samples collected in Shield (median: 0.31%, range: 0.06%-0.57%), this difference was not statistically significant (*p* = 0.12). Of reads classified as human, 1.2%-4.4% aligned to miRbase across the eight samples with no differences by preservative (*p* = 0.86). Most remaining human reads did not align uniquely (across the eight samples the percentage ranged from 81.3%-95.2%). For the remaining human reads that did align uniquely (4.8%-18.7% across the eight samples), the majority were classified as protein-coding or mitochondrial (**Additional File 3, Supplemental Table 1**).

A total of 42 human miRNAs were detectable across the eight samples based on a criterion of ≥10 raw counts in at least one sample (**Additional File 4, Supplemental Table 2**). Unsupervised hierarchical clustering of these miRNAs demonstrated strong clustering by child but not preservative (**Figure 4**), and normalized counts did not differ significantly between the two preservatives for any of the 42 detectable miRNAs after multiple testing correction (*p*_FDR_ ≥ 0.05).

**Figure 4:**
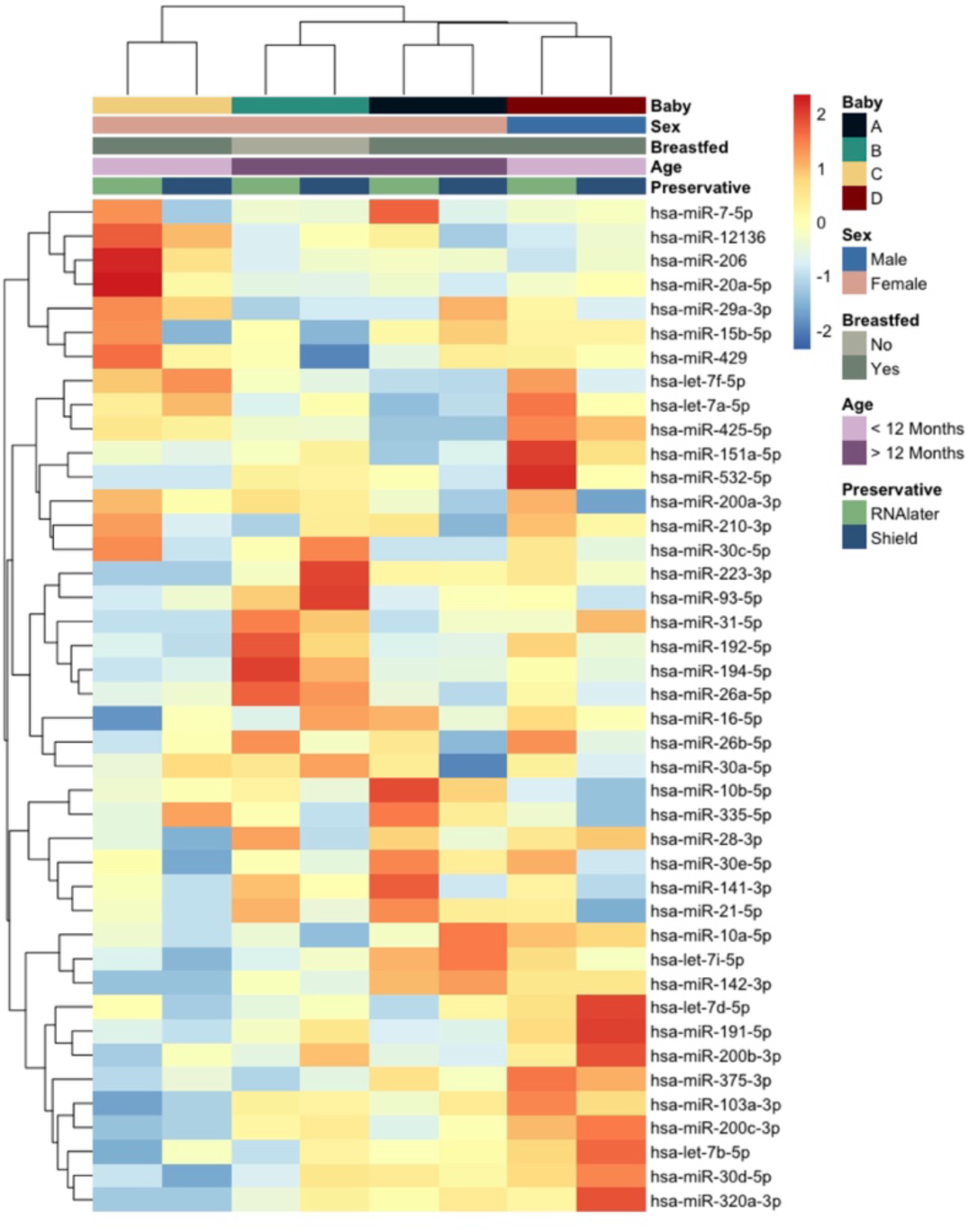
Scaled Heatmap of Detectable miRNAs in Infant Stool. Included miRNAs reflect the 42 miRNAs with 10 or more counts in at least one sample prior to normalization. The heatmap presents Z-score scaled counts for each miRNA. Manhattan distance was used for hierarchical clustering of both miRNAs and samples.

The top 10 most abundant miRNAs, ranked by normalized counts, are presented in **Table 2**.

**Table 2.** Summary of Top 10 miRNAs Ranked by Normalized Counts.

| Rank | miRNA | Length | Raw Counts |  | TMM Normalized |  |
| --- | --- | --- | --- | --- | --- | --- |
|  |  |  | Mean (SD) | Range | Mean (SD) | Range |
| 1 | hsa-miR-194-5p | 22 | 150 (170) | 11 - 443 | 5.9 (5.0) | 1.5 - 15.8 |
| 2 | hsa-miR-16-5p | 22 | 93 (82) | 21 - 284 | 4.2 (0.8) | 2.8 - 5.0 |
| 3 | hsa-miR-141-3p | 22 | 80 (73) | 20 - 230 | 3.5 (1.2) | 2.2 - 5.6 |
| 4 | hsa-miR-200c-3p | 23 | 71 (80) | 6 - 253 | 2.7 (1.5) | 0.8 - 4.9 |
| 5 | hsa-miR-21-5p | 22 | 58 (54) | 15 - 172 | 2.6 (0.8) | 1.2 - 3.7 |
| 6 | hsa-miR-191-5p | 23 | 62 (73) | 7 - 223 | 2.4 (1.7) | 0.8 - 5.9 |
| 7 | hsa-miR-26a-5p | 22 | 46 (42) | 12 - 129 | 2.0 (0.7) | 1.3 - 3.2 |
| 8 | hsa-let-7a-5p | 22 | 44 (56) | 16 - 180 | 1.9 (0.7) | 0.9 - 3.0 |
| 9 | hsa-let-7f-5p | 22 | 40 (55) | 9 - 173 | 1.7 (1.0) | 0.7 - 2.9 |
| 10 | hsa-miR-200a-3p | 22 | 31 (32) | 12 - 106 | 1.4 (0.4) | 0.8 - 1.8 |

### Gene Set Enrichment Analysis

MirDIP identified 1,340 unique high confidence predicted target genes for the top 10 most abundant miRNAs identified in infant/toddler stool. These predicted target genes were overrepresented (*p*_FDR_ <0.050) in 446 GO Biological Processes and 27 PANTHER pathways, with the top pathways being Regulation of DNA-Templated Transcription and the epidermal growth factor (EGF) Receptor Signaling Pathway, respectively (**Figure 5**).

**Figure 5:**
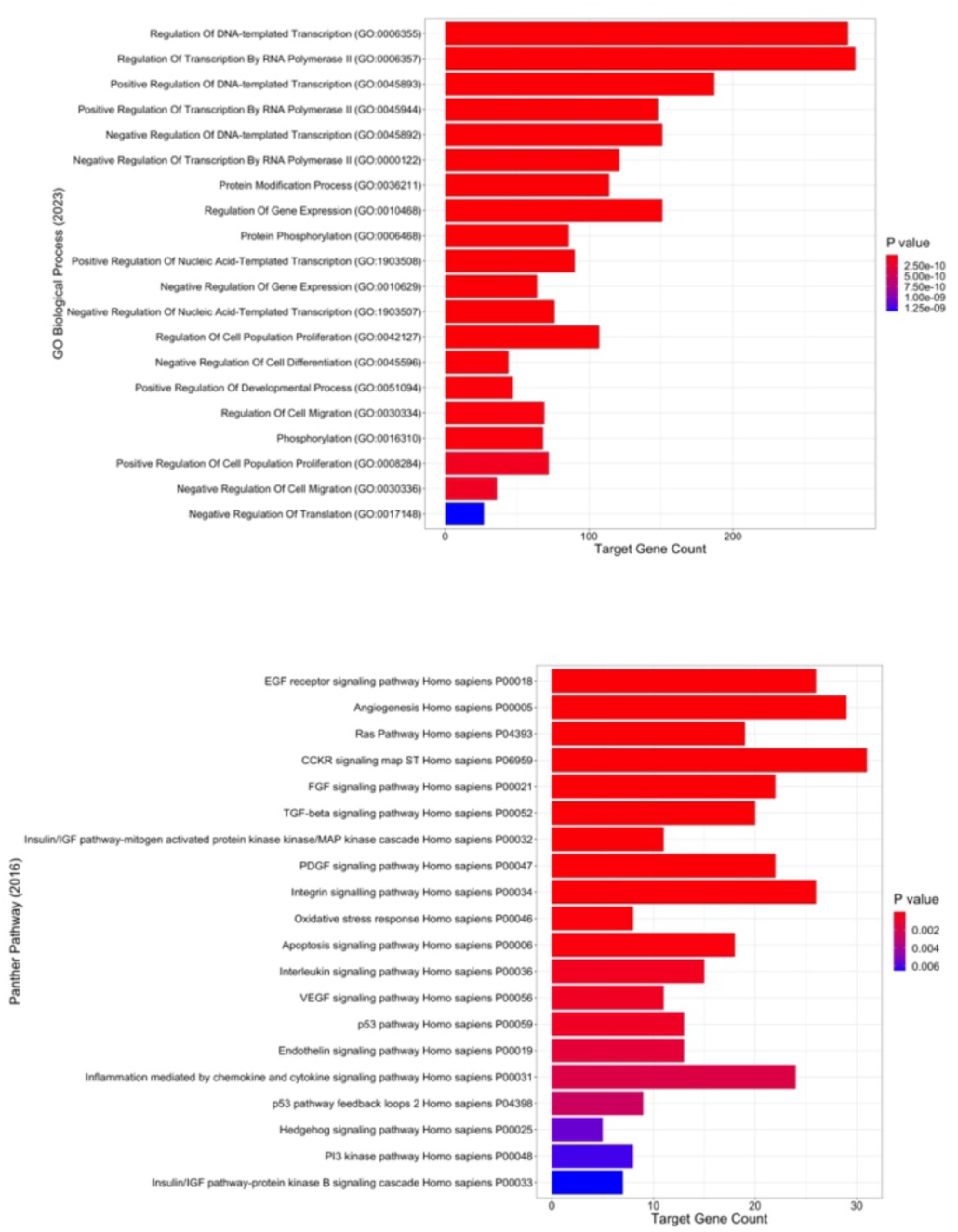
GSEA for Target Genes of the 10 Most Abundant miRNAs for (A) GO biological processes and (B) PANTHER pathways. Gene set enrichment analysis was conducted for 1,340 unique high confidence target genes, identified via mirDIP (version 5.2), of the top 10 most abundant miRNAs. Benjamini-Hochberg-adjusted p-values are presented.

## Discussion

MiRNAs in infant/toddler stool may provide unique insights into the mechanisms by which early factors influence children’s health, such as the developing microbiome and dietary patterns. The primary goal of this study was to identify optimal collection and extraction methods for obtaining high-quality RNA from infant/toddler stool for miRNA profiling. A secondary goal was to gain an initial understanding of the miRNA transcripts present in stool during this early life stage to inform future research on their potential to serve as non-invasive biomarkers for pediatric health. Of the kits and preservatives evaluated, one extraction kit (BIOMICs) and two preservation solutions (RNAlater and Shield) yielded high-quality RNA from infant/toddler stool, and no differences were observed in the levels of specific miRNA transcripts across the two preservatives. Using these collection and extraction protocols and a conservative approach for filtering out microbial reads, 42 unique mature human miRNA transcripts were identified in infant/toddler stool. A number of these miRNAs are known to be involved in the maintenance of gut barrier function (miR-194a-3p) [48], mucosal immunity (miR-200c-3p) [49], and cell proliferation (miR-26a-5p) [50, 51], which collectively contribute to gut homeostasis.

Over the last decade, 50 published studies have isolated RNA from adult stool samples. However, only nine studies have focused on stool from infants or young toddlers, and the majority either reported that they did not collect stool in a preservative [22–25, 30] or did not provide sufficient detail to determine if a preservative was used [26–29, 52]. Yet, use of a fixative that can preserve RNAs may be particularly important for stool which contains abundant amounts of RNases [35, 53]. This is an especially important consideration when studying a dual host-microbe environment, as microbial transcripts vastly outnumber human transcripts in stool, and degraded microbial RNA may have high sequence similarity to human miRNAs. Thus, degraded microbial RNA may be misclassified as human miRNAs. Minimizing RNA degradation is therefore important for downstream miRNA-seq analysis [34]. As expected, in the current study the highest degree of RNA degradation (based on RQN) was observed for infant/toddler stool samples that were collected without a preservative (median RQN: 2.5).

Currently, none of the nine published studies that specifically profiled miRNAs in infant/toddler stool reported use of an RNA preservation solution during stool collection, and the number of unique miRNA transcripts identified has varied substantially. For example, Carney *et al.* [23] used miRNA-seq for stool samples collected from infants 0-12 months of age to identify miRNAs associated with early growth and the gut microbiome and reported 53 detectable mature miRNAs. The full list of these miRNAs was not reported. However, 13 of the miRNAs that were associated with either infant growth or the gut microbiome overlapped with miRNAs that we also found to be detectable in infant/toddler stool. Furthermore, several of these common miRNAs were the most abundant miRNAs identified in our study (e.g., miR-194-5p, miR-16-5p, miR-141-3p, miR-200c-3p, and miR-30d-5p). In contrast, Kazokova, *et al.* [27] reported identifying 1,918 detectable miRNAs in meconium and infant stool samples using miRNA-seq. The much larger number of miRNAs reported in their study could be due to both the stool collection protocol used (no preservative) and the lack of a filtering step to remove microbial transcripts, which could have led to some reads being misclassified as human miRNAs, as well as library depth and threshold of reporting. Recent studies in adults that used a preservative (NPS) and RNA extraction kit (Norgen) that did not yield high quality RNA in our study have also reported much larger numbers of detectable human miRNA transcripts [38, 40, 41]. While this may be due in part to differences in life stage, misclassification of degraded microbial RNA could be a contributing factor, as no RNA integrity measures were reported in these studies, and microbial reads were not removed prior to human miRNA annotation.

Notably, all the preservative solutions evaluated here decreased RNA degradation compared to no preservative. However, RNAlater and Shield yielded the highest average RQN with median values of 9.8 and 9.4, respectively. High RQN values were observed despite potential lags between the sample being voided and the parent noticing the stool prior to transferring it to the respective preservative. Although samples collected in RNAlater generally yielded a higher percentage of reads being classified as human as compared with Shield, these differences were not statistically significant. The percentage of human reads aligning to miRbase also did not differ significantly by preservative, nor were any statistically significant differences observed between RNAlater and Shield for the abundance of specific miRNA transcripts. Furthermore, when comparing miRNA profiles of paired samples collected in RNAlater versus Shield, stronger clustering was observed by child than by preservative type. Collectively, these results suggest that while collecting infant/toddler stool in RNAlater may potentially improve sensitivity for downstream miRNA analysis, infant/toddler stool collected in RNAlater and Shield generally yields similar miRNA profiles.

As expected, the majority of reads in infant/toddler stool were microbial (>98% for each sample). This finding is consistent with two prior studies, one in infants [27] that did not use a preservative for stool collection, and one in adults [40] that collected stool in NPS. Of the reads that were classified as human, most did not align uniquely, and some aligned to coding and mitochondrial genes, likely reflecting degraded RNA. However, across the eight stool samples profiled a total of 42 unique mature miRNA transcripts were detectable. Many of the most abundant miRNAs in infant/toddler stool have been associated with biological processes relevant to gut homeostasis, such as epithelial cell differentiation and proliferation, epithelial-to-mesenchymal transition, maintenance of gut barrier function, and regulation of mucosal immunity. For example, the most abundant miRNA (miRNA-194) is known to be important for cell differentiation [48, 54] and is involved in epithelial-to-mesenchymal transition. A study of adults by Martinez, *et al.* [55] also reported that miRNA-16 is involved in dysregulation of intestinal immune system activation and epithelial barrier function, both of which contribute to the pathophysiology of irritable bowel syndrome. Additionally, Chen, *et al.* [56] showed that miR-141-3p can alleviate inflammatory response and oxidative damage in necrotizing enterocolitis through its regulation of inflammatory cytokines (IL-1β, IL-6, and TNF-α), and oxidative stress markers (MPO, MDA, and SOD). Moreover, Rawat, *et al.* has demonstrated that miR-200c-3p can alter the intestinal epithelial tight junction (TJ) barrier acting through the TJ protein occluding [49]. Finally, miR-21 and let-7 miRNAs have been shown to induce anti-inflammatory effects in response to Toll-like receptor mediated inflammatory responses [52, 57, 58]. Target genes of the top 10 most abundant miRNAs in infant/toddler stool were enriched in GO biological processes involved in a very broad range of cellular activities (e.g., DNA-templated transcription). However, several of the enriched PANTHER pathways, including EGF receptor, Ras, and MAP kinase signaling, play important roles in regulating intestinal development and intestinal barrier function. Thus, while a limited number of human miRNAs with generally low abundance were identified in infant/toddler stool, the specific miRNAs present may play important roles in intestinal development and gut homeostasis and have the potential to serve as useful biomarkers of child intestinal health.

Strengths of this study include comprehensive miRNA profiling using miRNA-seq and the collection of paired stool aliquots from the same diaper into two different preservatives, such that miRNA profiles could be directly compared across these two conditions. There are also several important considerations. First, it is possible that the approach used to distinguish human from microbial reads may have underestimated the true number of human miRNAs present in infant/toddler stool and their abundance. However, by using this conservative approach we were able to identify human miRNAs in infant/toddler stool with high confidence. This is important given the large quantities of microbial RNA present in stool that could be misclassified as human RNA when degraded. Notably, this study was designed to evaluate mature miRNAs given the growing interest in leveraging these as potential biomarkers of child health and development. However, pre-miRNAs and other classes of small RNAs have also been reported in stool [23, 27]. It will therefore be important to conduct future studies that compare these protocols for other classes of small RNAs. Finally, it is possible that this study was underpowered to quantify differences in levels of specific miRNA transcripts between samples collected in RNAlater versus the Shield preservative. There could be value in conducting future studies that collect a larger number of paired stool samples in these two preservatives to determine if collection in RNAlater improves the yield of low abundance miRNAs.

### Conclusions

Through this initial study, we identified two protocols that yield high-quality RNA for stool collected from infants and young toddlers. We also confirmed the presence of 42 mature human miRNAs in infant/toddler stool, many of which have known roles in gut homeostasis. This information can be used to inform the future collection of stool from young children to investigate the potential of these miRNAs to serve as non-invasive biomarkers for pediatric health.

## List of Abbreviations

EGF: epidermal growth factor
miRNA: microRNA
NP: No Preservative
NPS: Norgen Preservative Solution
Quick-RNA: Zymo Quick-RNA Fecal/Soil Microbe Microprep Kit
RQN: RNA Quality Numbe
Shield: Zymo DNA/RNA Shield Fecal Collection Tubes and Preservation Solution
TMM: Trimmed Mean of M-Values

## Declarations

### Ethics approval and consent to participate

All protocols were reviewed and approved by the Committee for the Protection of Human Subjects (CPHS), which is the Institutional Review Board for Dartmouth College. All participants provided written informed consent prior to participating.

### Consent for publication

This manuscript does not include any identifiable data from individual study participants. However, study participants consented to their use of their data for publications resulting from this research.

### Availability of data and materials

The datasets created and analyzed for this study are available from the corresponding author on reasonable request. However, requests for data access will require additional review by the Dartmouth College CPHS.

### Authors’ contributions

CGH conceptualized and designed the study; oversaw data acquisition, analysis, and interpretation of the data; and contributed to writing the initial draft of the paper. DAA contributed to the design of the study; data acquisition; interpretation of the data; and writing the initial draft of the paper. SMS contributed to data analysis and interpretation of the data and substantially revised the paper. MEM contributed to data analysis, interpretation of the data, and contributed to writing the initial draft of the paper. FWK contributed to data acquisition, interpretation of the data, and substantially revised the paper. HWT contributed to data acquisition. HEL contributed to the interpretation of the data and substantially revised the paper. AGH contributed to the interpretation of the data and substantially revised the paper. JG contributed to interpretation of the data and substantially revised the paper. BCC, JCM, ALH, and MRK substantially revised the paper.

