## Additional figures 1&2, Supplemental Tables 1&2 for "Optimizing Protocols for MicroRNA Profiling of Infant and Toddler Stool"

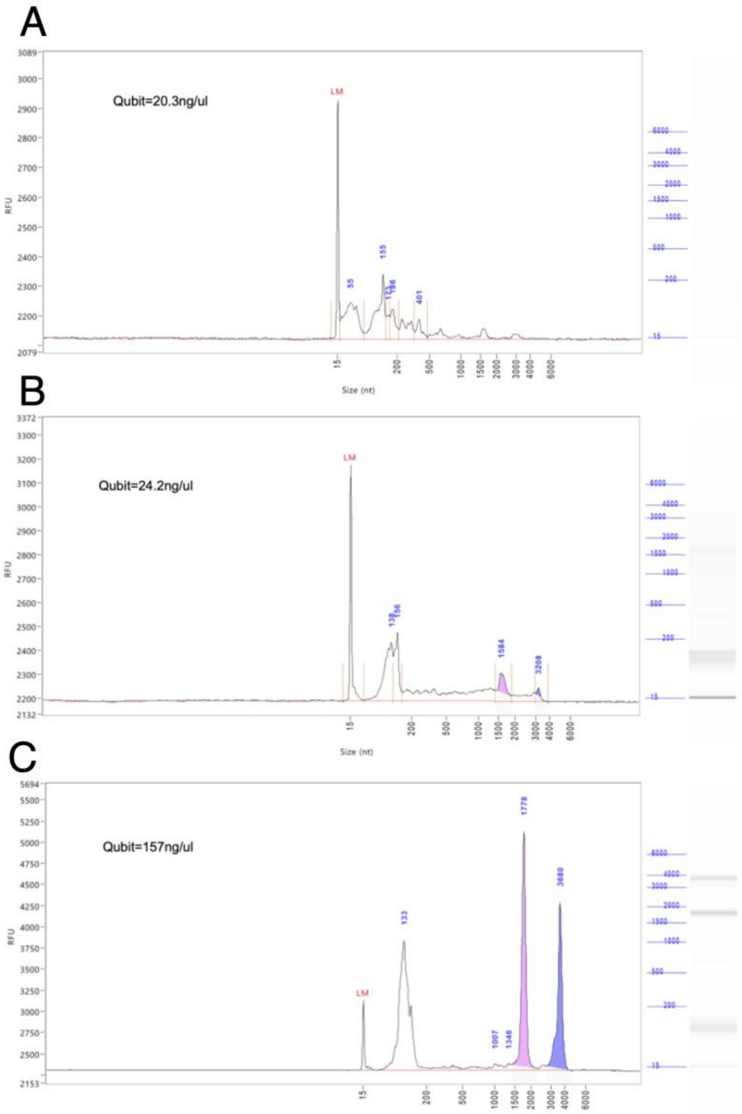


Additional File 1

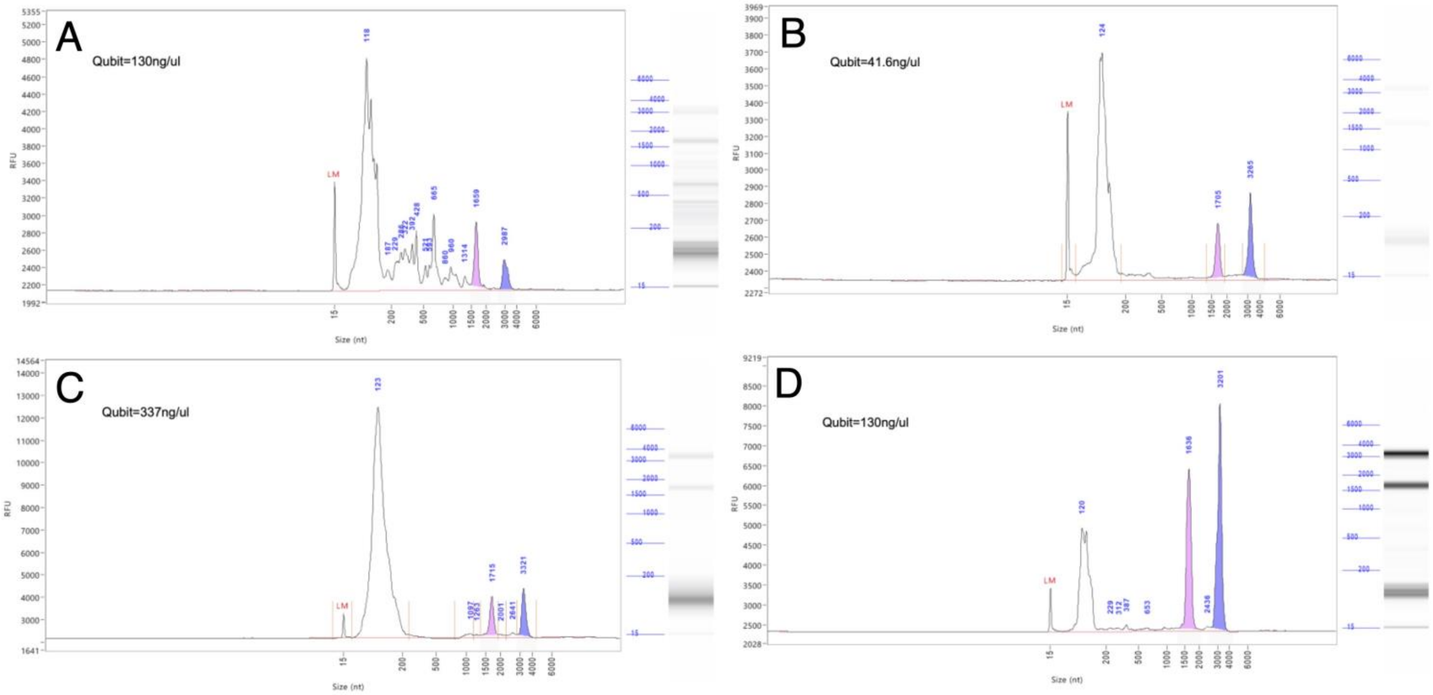


Additional File 2

**Supplemental Table 1.** Median (Range) percentage of uniquely aligning human reads that did not align to miRbase by RNA class for each preservative.

| lncRNA | | miscRNA | | MT | | Protein_coding | | snoRNA | | snRNA | |
|---|---|---|---|---|---|---|---|---|---|---|---|
| RNAlater | Shield | RNAlater | Shield | RNAlater | Shield | RNAlater | Shield | RNAlater | Shield | RNAlater | Shield |
| 0.18 (0.03-0.60) | 0.13 (0.04-0.72) | 0.01 (0.00-0.03) | 0.01 (0.00-0.04) | 0.21 (0.08-2.51) | 0.20 (0.11-2.00) | 0.42 (0.06-2.21) | 0.32 (0.09-2.08) | 0.03 (0.01-0.12) | 0.09 (0.00-0.15) | 0.01 (0.00-0.03) | 0.01 (0.00-0.05) |

RNA classes with a median value <0.01% not shown

**Supplemental Table 2.** Detectable miRNAs in Infant Stool from Four Infants Collected in RNAlater vs. Shield

| miRNA | All Samples Mean ± SD | All Samples Median (Range) | RNAlater Mean ± SD | RNAlater Median (Range) | Shield Mean ± SD | Shield Median (Range) | P | $P_{FDR}$ |
|---|---|---|---|---|---|---|---|---|
| miR-194-5p | 150 ± 170 | 62 (11-443) | 225 ± 221 | 224 (11-443) | 74 ± 61.4 | 54.5 (24-163) | 0.36 | 0.72 |
| miR-141-3p | 80 ± 73 | 46 (20-230) | 123 ± 84 | 119 (25-230) | 36 ± 13 | 37 (20-51) | 0.04 | 0.34 |
| miR-16-5p | 93 ± 82 | 76 (21-284) | 126 ± 112 | 100 (21-284) | 60 ± 20 | 63 (35-77) | 0.71 | 0.83 |
| miR-21-5p | 58 ± 54 | 36 (15-172) | 90 ± 64 | 84 (18-172) | 27 ± 10 | 28 (15-39) | 0.02 | 0.34 |
| miR-26a-5p | 46 ± 42 | 31 (12-129) | 66 ± 54 | 62 (12-129) | 26 ± 13 | 23 (17-43) | 0.17 | 0.56 |
| miR-200c-3p | 71 ± 80 | 43 (6-253) | 95 ± 110 | 60 (6-253) | 47 ± 35 | 43 (9-92) | 0.03 | 0.34 |
| miR-191-5p | 62 ± 73 | 36 (7-223) | 78 ± 99 | 40 (10-223) | 46 ± 46 | 34 (7-109) | 0.27 | 0.67 |
| miR-192-5p | 30 ± 35 | 15 (4-102) | 47 ± 46 | 40 (5-102) | 14 ± 9 | 13 (4-24) | 0.15 | 0.56 |
| miR-26b-5p | 30 ± 33 | 17 (6-105) | 47 ± 43 | 38 (6-105) | 13 ± 5 | 13 (8-18) | 0.20 | 0.56 |
| miR-200a-3p | 31 ± 32 | 19 (12-106) | 47 ± 41 | 35 (13-106) | 16 ± 5 | 14 (12-22) | 0.10 | 0.56 |
| let-7f-5p | 40 ± 55 | 19 (9-173) | 62 ± 75 | 31 (14-173) | 17 ± 7 | 18 (9-25) | 0.42 | 0.77 |
| miR-30d-5p | 33 ± 34 | 26 (3-110) | 44 ± 46 | 29 (6-110) | 23 ± 16 | 23 (3-42) | 0.62 | 0.81 |
| let-7a-5p | 44 ± 56 | 25 (16-180) | 63 ± 79 | 29 (16-180) | 25 ± 8 | 25 (16-35) | 0.93 | 0.98 |
| miR-10a-5p | 29 ± 27 | 23 (9-91) | 39 ± 36 | 29 (9-91) | 18 ± 8 | 18 (9-27) | 0.98 | 0.995 |
| miR-10b-5p | 16 ± 12 | 13 (3-35) | 24 ± 13 | 27 (5-35) | 9 ± 6 | 8 (3-17) | 0.18 | 0.56 |
| miR-30e-5p | 22 ± 25 | 12 (2-81) | 36 ± 32 | 27 (7-81) | 9 ± 5 | 10 (2-14) | 0.03 | 0.34 |
| miR-320a-3p | 26 ± 25 | 21 (1-77) | 31 ± 33 | 22 (1-77) | 22 ± 19 | 20 (1-48) | 0.15 | 0.56 |
| miR-103a-3p | 23 ± 28 | 14 (1-88) | 33 ± 38 | 21 (1-88) | 13 ± 7 | 14 (12-22) | 0.74 | 0.83 |
| miR-206 | 14 ± 4 | 13 (11-20) | 17 ± 3 | 17 (12-20) | 12 ± 2 | 12 (10-14) | 0.75 | 0.83 |
| let-7i-5p | 16 ± 17 | 11 (1-54) | 22 ± 22 | 16 (3-54) | 9 ± 6 | 10 (1-16) | 0.64 | 0.81 |
| let-7b-5p | 22 ± 24 | 14 (1-77) | 27 ± 34 | 15 (1-77) | 17 ± 11 | 14 (7-32) | 0.04 | 0.34 |
| miR-375-3p | 22 ± 32 | 9 (2-99) | 33 ± 45 | 15 (2-99) | 12 ± 9 | 9 (5-26) | >0.99 | >0.99 |
| miR-335-5p | 7 ± 5 | 5 (2-17) | 10 ± 6 | 10 (2-17) | 4 ± 1 | 4 (3-5) | 0.63 | 0.81 |
| miR-7-5p | 6 ± 5 | 5 (0-15) | 10 ± 5 | 10 (5-15) | 3 ± 2 | 3 (0-5) | 0.20 | 0.56 |
| miR-12136 | 7 ± 3 | 6 (1-11) | 9 ± 2 | 9 (6-11) | 5 ± 3 | 6 (1-6) | 0.62 | 0.81 |
| miR-93-5p | 7 ± 7 | 4 (1-19) | 9 ± 9 | 9 (1-19) | 5 ± 4 | 3 (2-11) | 0.51 | 0.81 |
| miR-210-3p | 9 ± 11 | 6 (1-36) | 14 ± 15 | 8 (4-36) | 5 ± 4 | 5 (1-8) | 0.41 | 0.77 |
| miR-28-3p | 5 ± 6 | 4 (0-16) | 8 ± 6 | 8 (1-16) | 2 ± 3 | 2 (0-6) | 0.17 | 0.56 |
| let-7d-5p | 12 ± 14 | 8 (1-44) | 15 ± 19 | 7 (3-44) | 10 ± 9 | 8 (1-22) | 0.51 | 0.81 |
| miR-429 | 7 ± 7 | 5 (1-22) | 10 ± 8 | 7 (4-22) | 4 ± 2 | 4 (1-6) | 0.35 | 0.72 |
| miR-20a-5p | 6 ± 4 | 6 (2-16) | 8 ± 5 | 6 (5-16) | 4 ± 2 | 3 (2-6) | 0.34 | 0.72 |
| miR-151a-5p | 8 ± 10 | 4 (2-33) | 12 ± 15 | 6 (2-33) | 4 ± 2 | 4 (2-7) | 0.75 | 0.83 |
| miR-200b-3p | 9 ± 10 | 5 (0-30) | 10 ± 14 | 5 (0-30) | 8 ± 7 | 7 (2-18) | 0.09 | 0.56 |
| miR-31-5p | 5 ± 5 | 4 (0-15) | 6 ± 7 | 5 (0-15) | 4 ± 4 | 4 (0-8) | 0.45 | 0.79 |
| miR-29a-3p | 6 ± 5 | 4 (3-18) | 8 ± 7 | 5 (3-18) | 4 ± 1 | 4 (3-5) | 0.79 | 0.85 |
| miR-30a-5p | 4 ± 4 | 3 (0-12) | 6 ± 5 | 5 (1-12) | 2 ± 2 | 2 (0-4) | 0.66 | 0.82 |
| miR-15b-5p | 3 ± 3 | 3 (0-10) | 5 ± 4 | 4 (2-10) | 2 ± 2 | 2 (0-5) | 0.33 | 0.72 |
| miR-142-3p | 3 ± 3 | 3 (0-10) | 4 ± 4 | 4 (0-10) | 2 ± 2 | 2 (0-3) | 0.63 | 0.81 |
| miR-223-3p | 3 ± 3 | 3 (0-11) | 4 ± 5 | 3 (0-11) | 2 ± 2 | 2 (0-5) | 0.61 | 0.81 |
| miR-30c-5p | 3 ± 3 | 2 (0-10) | 4 ± 4 | 3 (0-10) | 1 ± 2 | 1 (0-4) | 0.60 | 0.81 |
| miR-425-5p | 3 ± 4 | 1 (0-12) | 4 ± 6 | 2 (0-12) | 1 ± 1 | 1 (0-3) | 0.19 | 0.56 |
| miR-532-5p | 2 ± 4 | 1 (0-11) | 4 ± 5 | 2 (0-11) | 1 ± 1 | 1 (0-1) | 0.22 | 0.58 |
